# Chronic inflammation in ulcerative colitis causes long term changes in gobletcell function

**DOI:** 10.1101/2021.03.17.435871

**Authors:** Varsha Singh, Kelli Johnson, Jianyi Yin, Sunny Lee, Ruxian Lin, Helen Yu, Julie G. In, Jennifer Foulke-Abel, Nicholas Zachos

**Affiliations:** Division of Gastroenterology & Hepatology, Departments of Medicine, School of Medicine, Johns Hopkins University, Baltimore, MD 21205, USA; Department of Cellular and Molecular Physiology, Departments of Medicine, School of Medicine, Johns Hopkins University, Baltimore, MD 21205, USA; The University of Texas Southwestern Medical Center: Dallas, Texas 75390, US; Division of Gastroenterology and Hepatology, Department of Internal Medicine, University of New Mexico Health Science Center, Albuquerque, NM 87131, USA

**Author notes:** **Correspondence** Varsha Singh, PhD, Departments of Physiology and Medicine, Gastroenterology Division, Johns Hopkins University School of Medicine, Ross 933, 720 Rutland Avenue, Baltimore, MD 21205.

## Abstract

**Objective:** One of the features of ulcerative colitis (UC) is a defect in the protective mucus layer. This has been attributed to a reduced number of goblet cells (GC). However, it is not known whether abnormal GC mucus secretion also contributes to the reduced mucus layer. Our aims were to test the hypothesis that GC secretion was abnormal in UC with the changes persistent in colonoids even in the absence of immune cells.

**Design:** Colonoids were established from intestinal stem cells of healthy subjects (HS) and from patients with UC (inactive and active sites). Colonoids were maintained as undifferentiated (UD) or induced to differentiate (DF) and studied as 3D or monolayers on Transwell filters. Total RNA was extracted for quantitative real-time PCR analysis. Carbachol and PGE_2_ mediated stimulation followed by examination of mucus layer by MUC2 IF/confocal microscopy and TEM were performed.

**Results:** Colonoids derived from patients with UC can be propagated over many passages; however, they exhibit a reduced rate of growth and TEER compared with colonoids from HS. Differentiated UC colonoid monolayers form a thin and non-continuous mucus layer. UC colonoids have increased expression of secretory lineage markers: ATOH1 and SPDEF, including MUC2 positive GCs and ChgA positive enteroendocrine cells but failed to secrete mucin when exposed to the cholinergic agonist carbachol and PGE_2_, which caused increased secretion in HS. Exposure to TNF-α (5days), reduced the number of GC with a greater percentage decrease in UC colonoids compared to HS.

**Conclusions:** Abnormal mucus layer in UC is due to long term changes in epithelial cells that lead to abnormal mucus secretion as well as effects of the inflammatory environment to reduce the number of GC. This continued defect in GC mucus secretion may be involved in UC recurrence.

## INTRODUCTION

UC is a chronic relapsing colonic disorder. A frequent colonic abnormality in UC is a reduced mucus layer secreted by GCs. Mucus layer defects contribute to the UC pathophysiology by triggering immune responses and/or allowing increased and more proximate exposure to luminal bacteria, both of which can lead to further reduced barrier maintenance, mucosal damage, and defective absorption and increased fluid secretion. The mucus layer is secreted by GCs which primarily occur in differentiated colonocytes. Secretion of pro inflammatory cytokines in UC contribute to the destruction of epithelia barrier including mucus layer.^1 2^ However, even in the absence of endoscopic signs of active inflammation, the intestinal mucosa of UC patients in remission has a defective mucus layer and histological changes including branching of crypts, thickened muscularis mucosa, Paneth cell metaplasia, neuroendocrine cell hyperplasia. These changes suggest that the recovered intestine is permanently altered even after the inflammation has resolved. In fact, the intestinal epithelium of UC in remission has an expression profile that is significantly different from that found in healthy mucosa, that includes increases in expression of REG4, S100P, SERPINB5, DEFB1 and AQP3 and decreases in SLC16A1and AQP8 expression. Importantly these genes modulate epithelial cell growth, sensitivity to apoptosis and immune function.^3^ Other studies have shown that intestinal epithelium of IBD patients can harbor persistent alterations in gene expression or DNA methylation despite complete endoscopic and histologic remission.^2 4 5^ These changes could contribute to disease relapse which is common in UC.^3 6–8^ Altogether these results support the view that changes in the mucosa of patients with UC persist long after the inflammation has resolved.

We hypothesized that a long-term consequence of colonic inflammation is abnormal GC function that includes reduced stimulated mucus secretion. To test this hypothesis, we used an ex vivo human organoid/colonoid model from HS and from active and inactive mucosa of UC patients. Our results suggest that UC colonoids, that lack the presence of inflammatory cells, maintain an abnormal GC phenotype, with a reduced mucus layer due to defective cholinergic/PGE_2_ induced mucus secretion but with an increase in number of GC. Exposure of UC colonoids to TNF-α reduced the number of GCs, which occurred to a greater extent than in HS colonoids. Our results suggest that the abnormal mucus layer in UC is due to the effects of an active inflammatory environment to reduce the number of GCs as well as due to long-term changes in stimulated mucin secretion that persist even in the absence of inflammatory cells and exist in colonoids made from active and inactive UC.

## MATERIALS AND METHODS

### Patient population and biopsy collection

Colonic biopsies were obtained from HS (5) and UC patients (7) (Table 1) undergoing colonoscopies or from patients having colonic surgery for refractory UC. In all cases, informed consent was obtained using an experimental protocol approved by the Johns Hopkins University Institutional Review Board (IRB# NA_00038329). All procedures were performed in accordance with approved guidelines and regulations. Intestinal biopsies were collected from the ascending colon, descending colon, or sigmoid colon of HS screened with colonoscopy for colorectal cancer or gastrointestinal symptoms who had histologically normal colon. Seven UC patients had biopsies taken from area of uninvolved mucosa and/or active disease. Histologic status of the biopsies from colonoscopy or surgical samples are listed in **Table 1**. UC activity at the time of the colonoscopy was categorized according to the Mayo endoscopic subscore.^9^ Active UC was defined as a Mayo endoscopic subscore of ≥1; inactive disease was defined as a Mayo score of 0 in a previously involved segment. Colonoids were established via Hopkins Conte Basic and Translational Digestive Diseases Research Core Center Integrated Physiology Core (NIH/NIDDK P30).

**Table 1:** Clinical descriptions and origin of biopsies of non-IBD control (HS) and patients with Ulcerative colitis.

| Colon lines and disease | Number of subject included | Origin of biopsies |
| --- | --- | --- |
| UC inactive site | 5 | Ascending colon |
| UC inactive site | 2 | Sigmoid colon |
| UC active site | 2 | Descending colon |
| UC active site | 2 | Rectum |
| Healthy subject | 2 | Ascending colon |
| Healthy subject | 2 | Descending colon |
| Healthy subject | 1 | Sigmoid colon |

### Organoid culture and monolayer formation

Human colonoid cultures and monolayers were established utilizing the methods developed previously.^10 11^ Colonoids were maintained as cysts embedded in Matrigel (Corning #356231, USA) in 24-well plates and passaged as previously.^11^ Formation of enteroid monolayers was monitored by measurement of transepithelial electrical resistance (TEER). Undifferentiated 3D or monolayer cultures were induced to differentiate by removal of growth factors.^3^

### Quantitative Real-Time Polymerase Chain Reaction

Total RNA was extracted using the PureLink RNA Mini Kit (Life Technologies) according to the manufacturer’s protocol. Complementary DNA was synthesized from 1 to 2 μg of RNA using SuperScript VILO Master Mix (Life Technologies). Quantitative real-time polymerase chain reaction (qRT-PCR) was performed using Power SYBR Green Master Mix (Life Technologies) on a QuantStudio 12K Flex real-time PCR system (Applied Biosystems, Foster City, CA).

### Immunofluorescence staining, Confocal and TEM Image analysis

Analysis of MUC2 by immunofluorescence and confocal microscopy was carried out as previously reported using commercially available antibody.^5^ To evaluate qualitative mucin secretion colonoid monolayers were activated with the cholinergic analog carbachol (CCh) (25μM) to elevate intracellular Ca^2+^ and with PGE_2_ (1μM) to elevate cAMP in Kreb’s solution for 15mins followed with MUC2 staining and IF and TEM analysis.^12 13^ Primary antibody included Rabbit anti-MUC2 (Santa Cruz Biotechnology, USA; Cat#sc7314).

### Statistical analysis

Quantitative data are expressed as the mean ± and standard error of the mean (SEM). Statistical significance was determined using analysis of variance (ANOVA) with Bonferroni’s post-test (Prism GraphPad) to compare groups including a minimum n◻=◻3 replicates. A *p* ≤ 0.05 was considered statistically significant.

More detailed information is described in online supplementary materials and methods.

## RESULTS

### UC derived colonoids can be grown in culture over multiple passages; however, they exhibit a reduced growth rate

Human UC patient derived colonoids were propagated and compared with site matched HS colonoids. Similar to colonoids grown from HS, UC colonoids formed 3D spheroids and could be passaged at least 40 times. Figure 1A shows the phase contrast images of colonoids from HS and UC patients 5days post splitting. Morphologically, UC colonoids showed more budding structures compared with HS. However, when the growth of 3D spheroids was quantitated by measuring the number of spheroids per well after each split over time and for multiple passages, active disease UC colonoids grew slowly and formed less spheroids compared to inactive UC and HS (figure 1B). We have demonstrated that human colonoids can be grown as 2D monolayers.^14^ The progress of monolayer formation was monitored on a daily basis by a steady increase in transepithelial electrical resistance (TEER) (figure 1C). The monolayers were maintained in the undifferentiated (UD) crypt like state by growth in Wnt3A, RSPO1 and Noggin, while withdrawal of growth factors (Wnt3A and RSPO1) drove differentiation (DF) by 5days. As shown in figure 1C, active and inactive UC colonoid were delayed in establishing confluency and had lower TEER (Inactive UC: UD 700 Ω·cm^2^ ± 60, DF 1500 Ω·cm^2^ ± 60; n=10, *p*≤*0.05* vs HS; active UC: UD 600 Ω·cm^2^ ± 60, DF 1200 Ω·cm^2^ ± 80;n=10, *p*≤*0.05* vs HS) compared to monolayers from HS (UD 1200 Ω·cm^2^ ± 55, DF 2500 Ω·cm^2^ ± 55; n=10) measured at post plating day10 for UD and day 15 for DF (figure 1D).The slow growth of colonoids in 3D and 2D monolayer formation suggests that there are sustained differences within the epithelial stem cell compartment of the UC vs HS mucosa.

**Figure 1:**
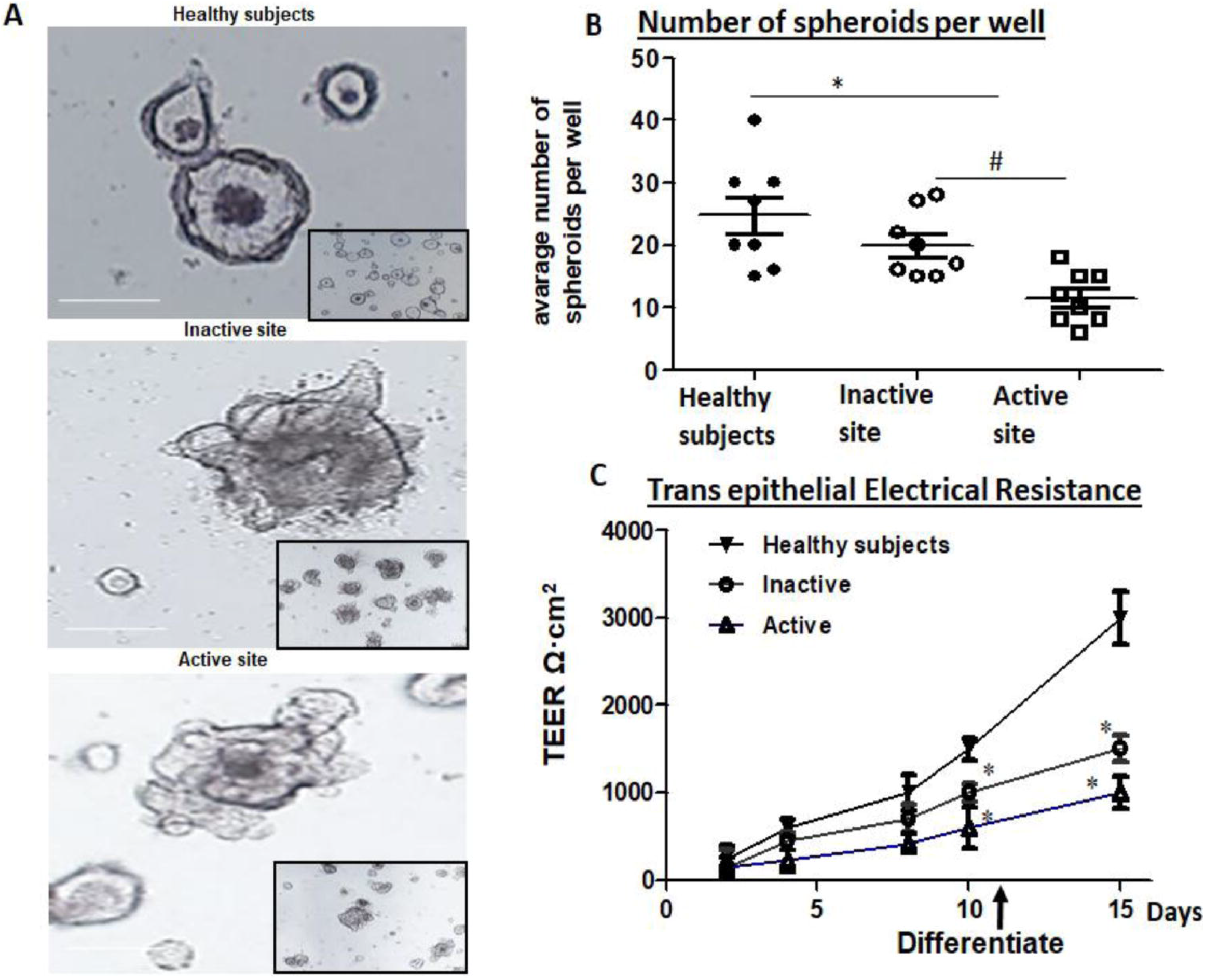
UC colonoids have differences in growth compared to colonoids from HS: A) A representative bright field image of 3D colonoid from HS, inactive and active site of UC patients. B) Number of 3D spheroids per well from HS, inactive and active site of UC patients. Quantitation of spheroids were made 2 days post splitting. C) Changes in TEER of colonoid monolayers rom HS (black triangle), inactive (circle), and active (triangle) UC. TEER of monolayer increased further upon 5 days of differentiation (Wnt3A and Rspond removal). Spheroids and monolayers from all the subjects were analyzed at least 3 times (n=5 HS, 4=UC active and 7=UC inactive), \**p* < 0.05 vs HS, ^#^*p* < 0.05 vs inactive UC. Scale bar 20μm.

### Colonoids derived from UC tissue form thin mucus layer and have defective barrier integrity

Active UC tissue have a reduced mucus layer and many UC colons have a reduced number of GC^14^. Similarly, colonoid monolayers made from the tissue derived from either inactive or active site of UC lacked a uniform mucus layer; instead they have a thin and a non-uniform mucus layer (figure 2A). We further analyzed the number of GC in these monolayers by counting MUC2 positive cells per monolayer. Surprisingly, differentiated UC monolayers from both active and inactive sites had a significantly higher number of GC compared with monolayers from HS (figure 2B). The primary component of mucus layer is MUC2, an extensively O-glycosylated molecule that forms polymeric sheets to which luminal bacteria attach and which provides a food source for the microbiota.^4 15^ O-glycans contribute to about 80% of its mass and therefore are important determinant of mucus properties. O-glycosylation of MUC2 occurs post-translationally in the Golgi apparatus. The primary enzymes in this process are the core 1 β1,3-galactosyltransferase (C1galt1), core 2 β1, 6*N-*acetylglucosaminyltransferases (C2GnTs) and core 3 β1,3-N-acetylglucosaminyltransferase (C3GnT).^16^ The mRNA levels of several enzymes responsible for glycosylation of mucin dimers were measure including C1galt1, C2GnT and C3GnT. Of the enzymes tested, C2GNT2, did not increase with differentiation of UC colonoids as it occurred in HS colonoids. Similar results were seen in colonoids from inactive and active sites of UC patients (figure 2C). In contrast, mRNAs of C1galt1 and C3GnT were not significantly different from HS (data not shown).

**Figure 2:**
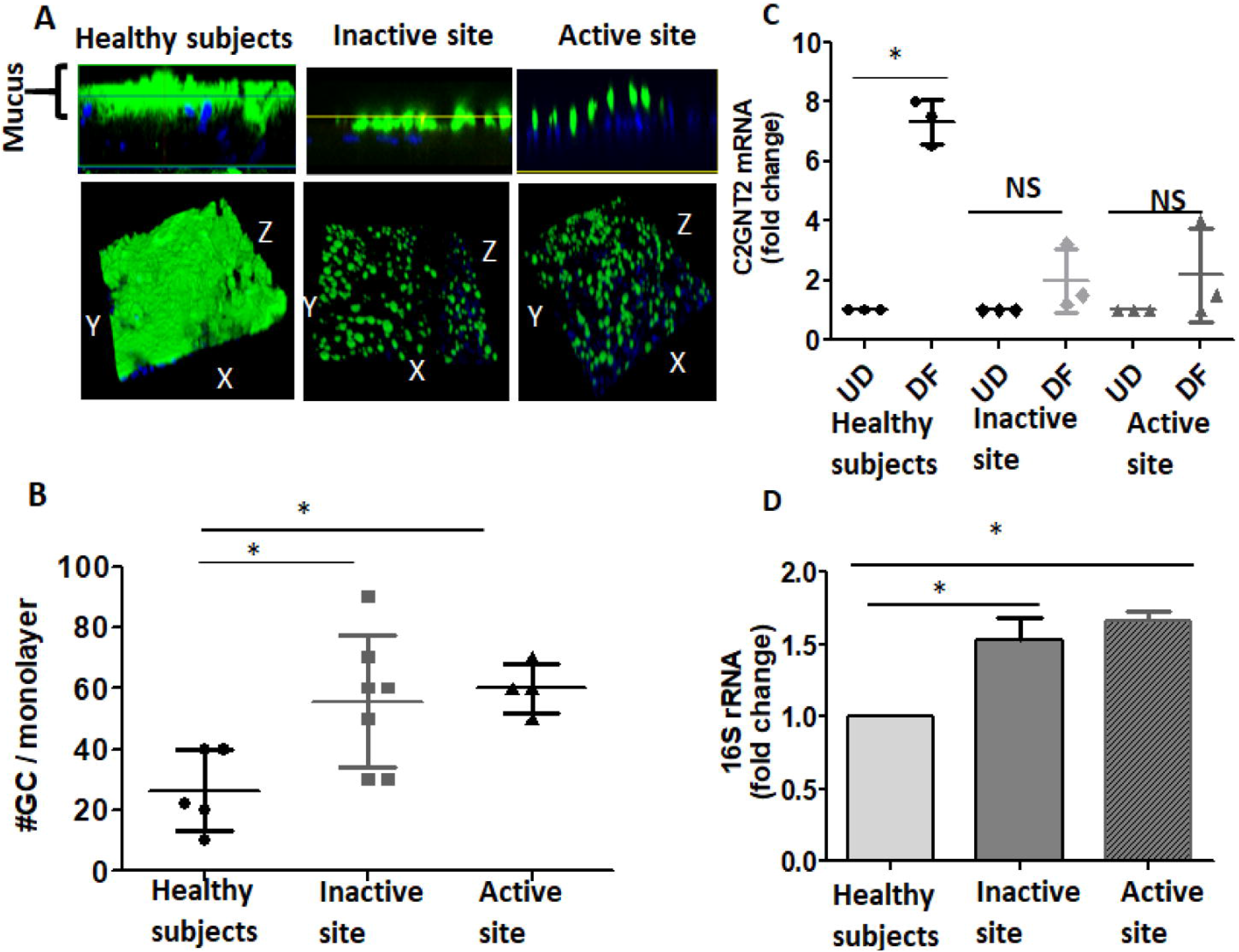
UC colonoids have defects in mucus secretion and barrier function: A) Methanol–Carnoy’s fixed differentiated colonoid monolayers stained with MUC2 (green), nucleus (blue). Representative confocal XZ (above) and 3D-XYZ (below) projections depicting the MUC2 layer in colonoids monolayer is shown. B) Average number of GC expressed post 5 days of differentiation of colonoid monolayers. C) Differences in the mRNA expression of C2GNT2 mRNA, post differentiation of monolayers from HS, inactive and active UC sites. D) Bacterial 16S rRNA expression in colonoids post 8h infection of differentiated monolayers. A and B: multiple areas of monolayers from each group were analyzed (n=5 HS, 4=UC active and 7=UC inactive). C and D: n=3 monolayers from each group was analyzed at different times. Results are shown a Mean±SEMs. *p<0.05 vs HS. Scale bar 20μm.

We further investigated the barrier integrity by exposing differentiated colonoid monolayers to apical *Escherichia coli* (1×10^6^cfu/ml) (8h) and performed16S bacterial rRNA based real-time PCR analysis on total RNA extracted from monolayers. An increased amount of bacterial 16S-rRNA was present in monolayers from UC patients (inactive and active) as compared to HS, suggesting that UC colonoids have a defective mucus barrier (figure 2D).

### Activation of secretory lineage differentiation in UC compared with non-IBD controls

In order to investigate the differentiation status and GC related gene expression in UC colonoids, we performed qPCR expression analysis of a selected panel of genes in UD and DF colonoids from HS and UC patients. The expression of the stem cell gene, Lgr5 and cell proliferation marker, Ki67 were slightly but not significantly increased in both inactive and active UC colonoids as compared with HS (figure 3A). Nonetheless, the expression of both Lgr5 and Ki67 decreased with DF of UC colonoids as in HS. In addition, the expression of genes associated with mucus producing GC was determined. Shown in figure 3B, is a transcription factor atonal homolog 1 (ATOH1) which is a gatekeeper that controls the fate of intestinal progenitors. Intestinal progenitors with reduced Notch activity express high levels of ATOH1 and commit to a secretory lineage fate (figure 3B). Therefore, ATOH1 expression in UD and DF colonoids was measured. Both active and inactive UC colonoids in UD as well as DF states had significantly higher expression of ATOH1 compared with HS (figure 3C, left). The expression of transcription factors downstream of ATOH1 were also analyzed, including SPDEF and Ngn3 which specify differentiation and maturation of GC and enteroendocrine cells respectively (figure 3B). Similar to ATOH1, the expression of SPDEF and Ngn3 were significantly higher in both active and inactive UC colonoids compared with HS (figure 3C, middle, right respectively). The expression of MUC2 (GC marker), ChgA (enteroendocrine cells) and Lyz (Paneth cells) were also determined (figure 3D). MUC2 message was increased in the UD colonoids from both active and inactive UC compared to HS, while the message was not significantly different between DF colonoids from each group. In contrast, ChgA transcripts followed a pattern of upregulation in UD as well as in DF UC colonoids from active and inactive UC compared to HS. Lysozyme transcripts were increased in only some of the UC colonoids, but there was no consistent change compared to HS in UD or DF active or inactive UC colonoids.

**Figure 3:**
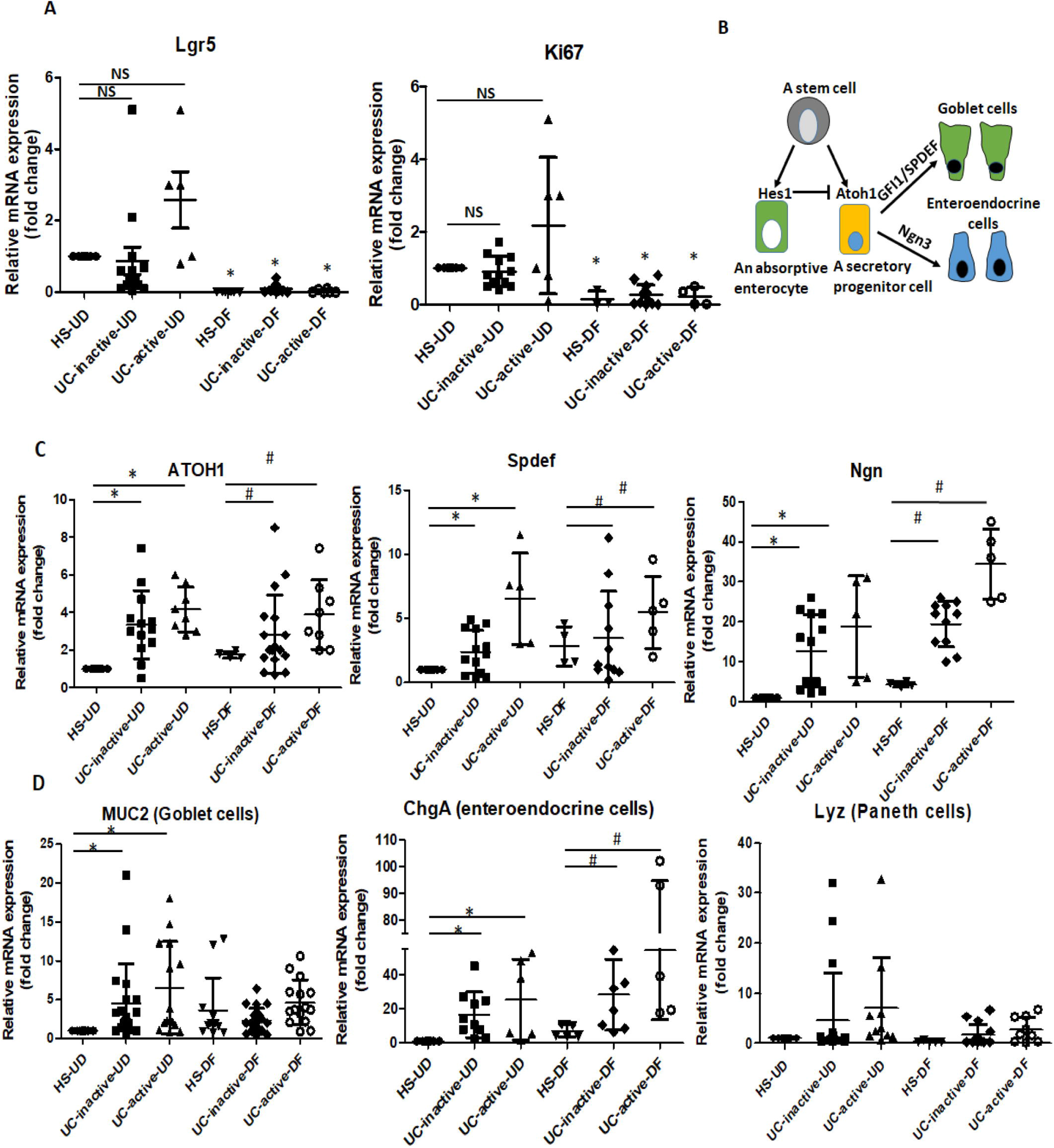
Differential gene expression profiles in UD and DF colonoids from HS compared with UC patients: Relative mRNA levels of: A) proliferation genes, B) schematic representation of absorptive and secretory pathways starting from a progenitor and the genes involved in this process, C) secretory lineage genes, D) genes specific to different cell types, by qPCR. Messenger RNA levels are normalized to *18S ribosomal RNA* expression. Result is normalized to HS set as 1 and expressed as fold change. Results are Mean±SEMs. *p<0.05 vs HS-UD; ^#^p<0.05 vs HS-DF; 3D colonoids from n=5HS, 4=UC active and 7=UC inactive sites; some colonoid lines were studied multiple times.

### UC colonoids differentially express ion transport proteins as compared with HS

To further define the differentiation states of UC colonoids, mRNA expression of several ion transport proteins and a carbonic anhydrase isoform was determined. These ion transporters and carbonic anhydrase isoform are known to play important roles in Cl^−^ and HCO_3_^−^ secretion, electroneutral Na^+^ absorption, and intracellular pH regulation under physiological and pathophysiological conditions and have been shown to undergo changes in expression with differentiation in intestinal epithelial cells.^17^ As reported previously and shown in figure 4, several ion transporters and carbonic anhydrase isoforms were up-regulated significantly at the mRNA level upon differentiation in HS colonoids. These included sodium hydrogen exchanger-3 (*NHE3)* (18.4-fold), *DRA* (13.6-fold), *CA2* (2.0-fold), *NHE1* (2.7-fold). In contrast, several ion transporters were down-regulated significantly after differentiation, including *NKCC1* (20.1-fold), potassium channel, voltage gated, subfamily E, regulatory subunit 3 (*KCNE3*) (4.2-fold), and *CFTR* (12-fold). In contrast, UC colonoids exhibited somewhat different mRNA expression patterns compared with HS. In the UD state, UC colonoids (inactive and active site) had significantly higher expression of NHE3 (inactive 27-fold; active 3.4fold), DRA (inactive 5-fold; active 3.2fold) and CA2 (inactive 2.1-fold; active 1.2-fold), and lower expression of CFTR (inactive 0.5-folds; active 0.3folds). Differentiation failed to cause significant change in the expression pattern of NHE3 and DRA. Importantly, when compared with DF HS, DF UC colonoids had significantly lower expression of NHE3 (inactive 2-fold; active 4-fold) and DRA (inactive 5.5-fold; active 8-fold). The mRNA levels of several other transporters were not significantly different between the groups: anion exchanger 2 (*AE2*), electroneutral Na^+^/HCO_3_^−^ co-transporter 1 (*NBCe1*), *NHE2* and putative anion transporter 1 (*PAT-1).* Overall this suggests that in undifferentiated conditions UC colonoids were partially differentiation based on the increased mRNA expression pattern of NHE3, DRA, CA-II and decreased CFTR expression. In contrast, in differentiated colonoids from inactive and active UC, there was no further or even reduced differentiation based on reduced NHE3, DRA, and slight increase (not significant) in NKCC1 expression. These data suggest that the pattern of differentiation and expression of multiple ion transporters and a carbonic anhydrase isoform in UC colonoids is different from HS.

**Figure 4:**
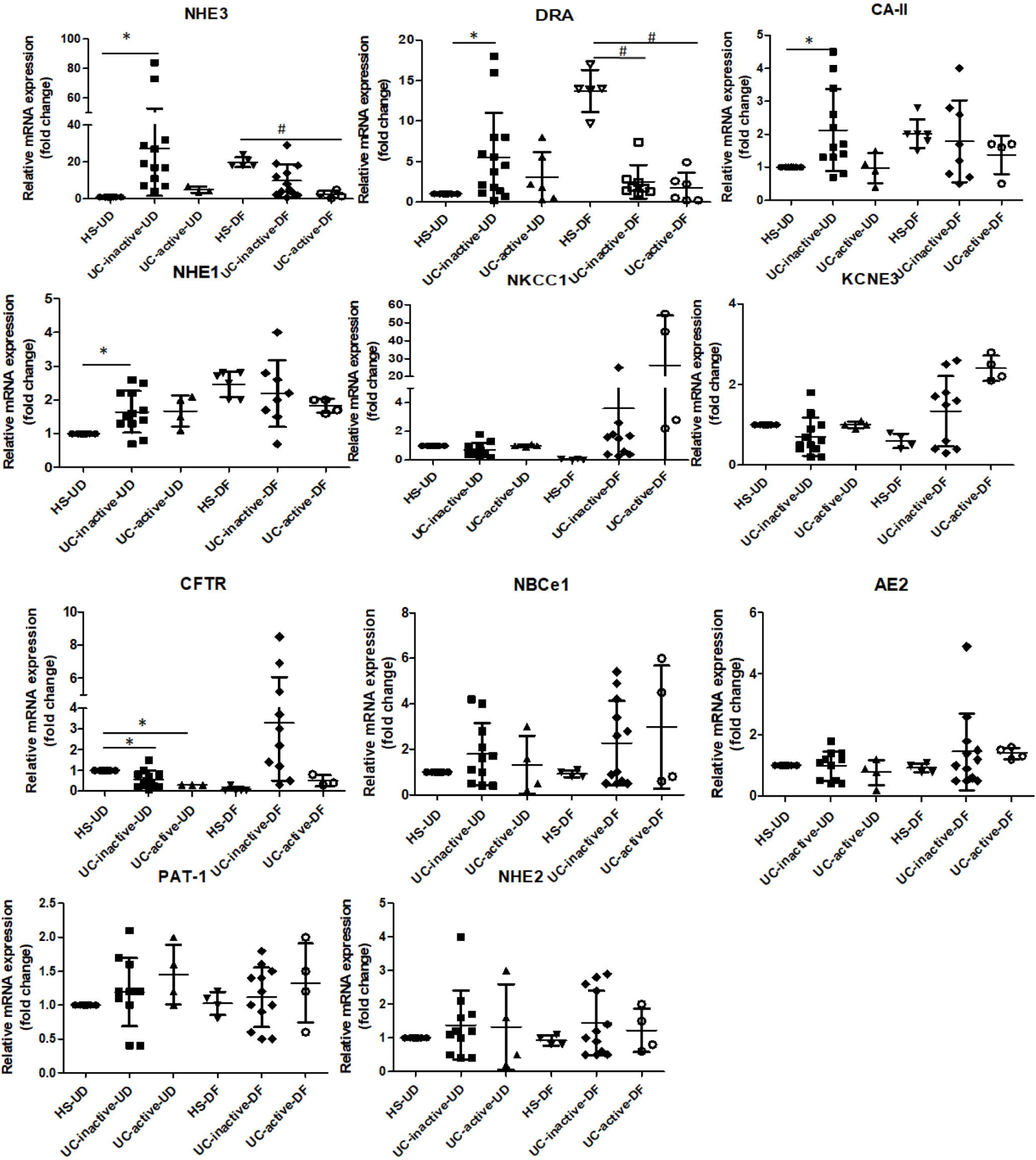
mRNA levels of selected ion transporters and carbonic anhydrase in UC colonoids compared with HS. A) The mRNA levels of selected ion transporters were determined by qRT-PCR and relative fold change between undifferentiated (UD) and differentiated (DF) colonoids were calculated using *18S ribosomal RNA* as endogenous control. Results are normalized to HS set as 1 and expressed as fold change. Mean±SEMs.*p<0.05 vs HS-UD; ^#^p<0.05 vs HS-DF; 3D colonoids from n=5-HS, 4=UC active and 7=UC inactive sites; some colonoid lines were studied multiple times.

### Goblet cells in UC colonoids do not respond to carbachol (Cch) and PGE_2_ mediated mucin secretion

In addition to synthesizing MUC2, goblet cells release stored MUC2 granules in response to cholinergic plus cAMP related stimuli. Multiple studies have found that Ca^2+^ signaling is required for the release of mucin-filled vesicles.^18 19^ In accordance with the known muscarinic cholinergic signaling pathway for mucin secretion, we treated UC monolayers with carbachol (Cch) (25μM) to elevate intracellular Ca^2+^ and with PGE_2_ (1μM) to elevate cAMP^20^. In contrast to Cch/PGE_2_ induced mucin secretion and creation of a thick mucus layer in colonoids from healthy subjects, monolayers from both inactive and active UC did not respond to the treatment (figure 5A). At the ultrastructure (TEM) level, GCs in HS had the expected appearance of granule-filled vesicles located just apical to the nucleus (figure 5B). Following stimulation with Cch+ PGE_2_, most of the GC in HS exhibited cavitation at the apical side. In contrast, GC in UC monolayers (both inactive and active) did not show any decrease in the mucin vesicles in response to the treatment. Overall, these results suggest that colonoids in UC can differentiate to GC, but have a compromised secretory function in response to cholingeric/cAMP stimulation.

**Figure 5:**
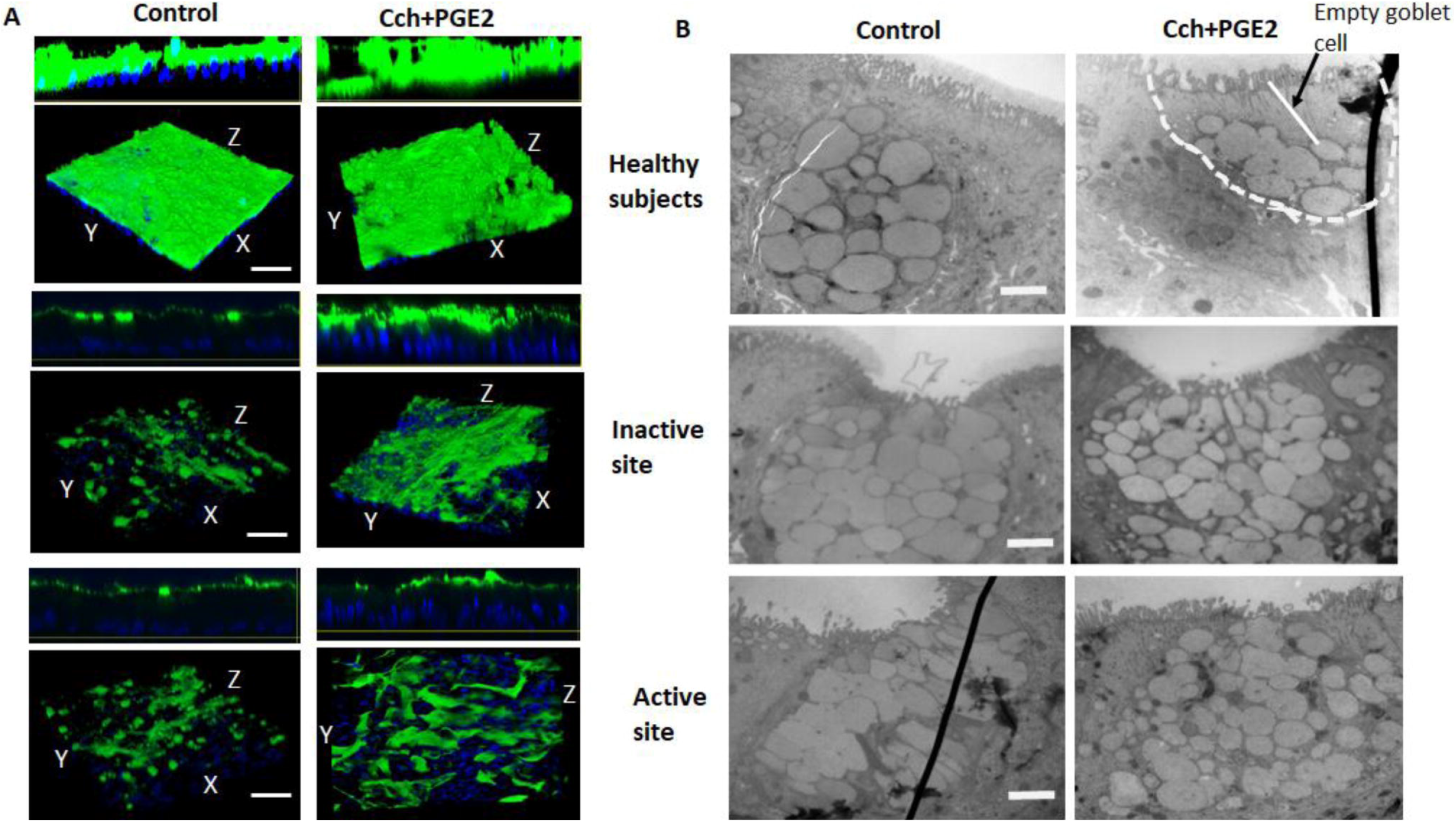
UC colonoids have defects in mucus secretion: Colonoid monolayers from HS, inactive and active UC sites were treated with carbachol (25μM) + PGE_2_ (1μM) for 15mins and then analyzed. A representative image from each group is shown. A) Methanol–Carnoy’s fixed monolayer stained for mucus layer, Muc2 (green), nucleus (blue). Scale bar 20μm. B) TEM of GC from control and Cch/PGE_2_ treated monolayers from different groups. Note the empty area on the apical side of GC in HS, treated with Cch/PGE_2_, but not in UC. n=3 monolayers from different subjects in each group. Scale bar 500nm.

### TNF-α treatment reduces GC number

Since the colonoid model is devoid of any immune cells, we hypothesized that the differences in GC number in our model from those reported in UC patient tissue samples is because of the absence of active inflammatory cytokines secreted by immune cells in UC patients. To test this hypothesis, we differentiated monolayers from HS and UC patients in presence of inflammatory cytokine TNF-α (5ng/ml, added freshly with media change at 2^nd^ day of 5day DF), followed by analysis of MUC-2 positive GC per monolayer. A representative example is shown in figure 6A and quantitation of multiple monolayers are shown in figure 6B, TNF-α treated monolayers had decreased numbers of MUC-2 positive GC in both HS (control: 40±12; TNF-α: 22±5.6) and UC colonoids (inactive control: 65±15; TNF-α: 20±12, active control: 71±14; TNF-α: 26±10,). The percent change in number of GC in UC was higher than in HS subject (UC inactive 69%; UC active 63%; HS: 45%). These results suggest that the decrease in GC number in UC patient tissue samples is dependent on active inflammatory cytokines.

**Figure 6:**
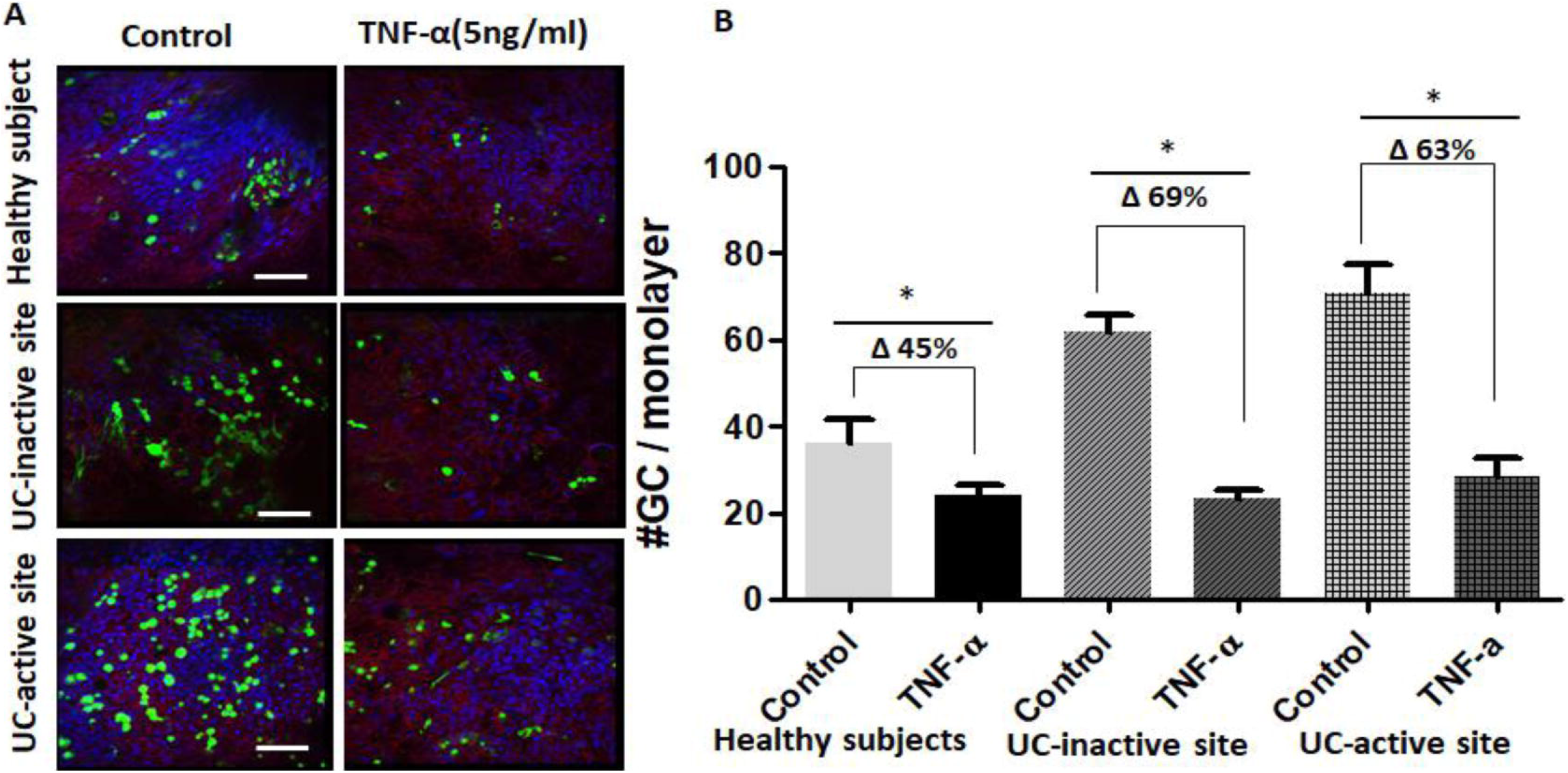
TNF-α (5ng/ml, 5 days) treatment decreases GC number: A) Monolayer from HS and UC- inactive and active sites were differentiated alone or with TNFα (5ng/ml) for 5days and MUC2 positive GC (green) were analyzed using confocal imaging. Fresh TNFα (5ng/ml) was added during media change, at second day of 5day period. B) Average number of GC expressed in untreated or TNFα treated monolayers. Results are Mean±SEMs.*p<0.05 vs control/untreated monolayers, n=3 separate monolayer from each group. Scale bar 20μm.

## DISCUSSION

In this study we provide a new mechanistic insight into the basis for the reduced mucus layer that is part of pathophysiology of UC. Although, a reduced number of GC as reported in many UC cases is considered as the sole cause of the reduced mucus layer, our studies suggest that the reduced mucus layer seen in UC patients is related to both reduced number and reduced secretory function of the remaining GC. Furthermore, we also provide evidence that the epithelial compartment in UC undergo alterations and have reduced expression of bicarbonate transporters: DRA and CFTR. Reduction in luminal HCO_3_^−^ is known to contribute to failure of the mucin to unfold. This is one of the components important in the multicomponent pathophysiology of UC.

Altered characteristics of epithelial cells in UC is thought to be largely due to the inflammatory environment. However, it was not known which of these changes revert back to normal once the inflammation is removed or whether some of them are imprinted in the epithelial compartment. In the present study, we took advantage of the ability to establish stem cell derived colonoids from active and inactive areas of UC that could be passaged at least 40 times and studied them in both the UD crypt like and DF upper crypt and surface cell state to begin defining some of these long term changes. Colonoids made from active and inactive areas of UC had properties distinct from colonoids made from the same colonic segments from healthy control subjects; for instance, the growth rate was much slower in colonoids from active UC and the TEER was significantly reduced in colonoids from both active and inactive UC. The reduced TEER is an indication of abnormal tight junctions and intestinal barrier function and duplicates a feature known to be present in patients with UC. Undifferentiated active and inactive UC colonoids had increased mRNA expression of proteins normally present in differentiated colonocytes, including NHE3, DRA, CA-II but had reduced expression of CFTR which is usually more highly expressed in the crypt; moreover, when the colonoids were exposed to conditions that led to differentiation in colonoids from HS, these genes either failed to increase or decreased in expression. MUC2 also behaved similarly and in a distinctly abnormal pattern, being increased in UD active and inactive UC colonoids, while there was no further increase with application of differentiation conditions. Decreased expression of DRA is reported in various inflammatory diarrhea and in UC patients^21^. Similarly, decreased mRNA expression of CFTR in UC colonoids is in accordance with the reports from animal model of colitis as well as from UC patients (DOI 10.21203/rs.3.rs-22104/v1-preprint). These results are consistent with long term effect of inflammation in UC colonoids exhibiting early differentiation, that fits with the reduced proliferation shown in figure 1B. However, the mechanisms for these long-term changes has not been identified.

The UC colonoids had an increased number of GC compared to HS colonoids. This was consistent with the increased level of ATOH1, a transcription factor that increases stem cell differentiation towards the secretory pathway. Moreover, there was also an increase in Chromogranin A positive enteroendocrine cells in DF UC colonoids, another part of the ATOH1 driven secretory cell developmental pathway. Several studies have reported greater numbers of enteroendocrine cells in the colonic mucosa of the patients with active UC, indicating similarity between the UC colonoid model and intact colon.^22 23^ In spite of higher expression of GCs, both active and inactive UC colonoids formed a thin mucus layer, suggesting defects at the level of the signaling pathways or secretory machinery required for mucus secretion.

Secretion of mucin release from GC was examined by exposure to the muscarinic agonist carbachol plus the cAMP agonist PGE_2_, agonists known to cause mucin exocytosis.^13^ Formation of a functional mucus layer is a result of a complex multi-step process. It starts with an increase in intracellular Ca^2+^ in response to activation of muscarinic M3 receptors, which is followed by fusion of mucin containing vesicles, compound exocytosis and finally mucin unfolding via HCO_3_^−^ exposure. Mucin release was markedly reduced in both active and inactive UC colonoids based on the measurement of changes in the mucus layer in colonoid monolayers and by examining the apical area of GC by TEM (figure 5). Further studies are required to understand which of the multiple steps in mucus secretion is abnormal in UC. The second contributor to a thin mucin layer in UC colonoids is related to dependence of mucus unfolding on luminal HCO_3_^−^. Although it is not known if the HCO_3_^−^ comes from adjacent epithelial cells or is more closely associated with the GC, but the mRNAs of both DRA and CFTR: two major colonic apical HCO_3_^−^ transporters were significantly reduced in the differentiated UC colonoids. The third likely contributor to abnormal GC mucin secretion is abnormal expression of C2GnT2, an enzyme responsible for O-glycosylation of MUC2. C2GnT2 is highly expressed in the mouse small intestine and colon and C2GnT2 deficiency reduces levels of core 2 and 4 O-glycans, as well as I-branching. Moreover, C2GnT2−/− mice exhibit increased susceptibility to DSS-induced colitis. Additional studies are required to determine if mucin glycosylation is abnormal in UC colonoids and to define the consequence of altered glycosylation on mucus layer formation.

A thin and defective mucus layer is a signature of UC and this has been attributed to the reduced number of colonic GC. In contrast, in our studies, UC colonoids had an increased number of GC compared to HS colonoids. One of the limitations of studies with stem cell derived intestinal organoids is that they only contain epithelial cells and lack the many additional cell types present in the normal intestine, including inflammatory and immune cells. Consequently, disease models using colonoids do not entirely duplicate the inflammatory or immune environment that plays a critical role in pathophysiology of many GI diseases, including IBD. To deal with this limitation, co-culture with additional cell types has been developed, while use of iPSC derived organoids includes some of the additional mesenchymal cells present in the colon. Based on this limitation, we hypothesized that the difference in GC number in UC tissue compared to UC colonoids might be due to lack of the inflammatory environment. In fact, when colonoids were exposed to TNF-α for 5days, there was a decrease in GC number in both HS and UC colonoids, with the reduction in number in the UC colonoids exceeding that in HS. This finding supports the interpretation that the reduced number of GC in UC is at least in part due to the local inflammatory environment. The conclusion from these studies is that the reduced protective mucus layer in UC is a consequence of both a reduced number of GC, which appears to be a reversible inflammation dependent phenomenon, and reduced mucin secretion by the remaining GC, which appears to be a long term part of the disease.

The current observations, further support several recent studies that have suggested that epithelial cells from the involved colonic mucosa of patients with UC acquire a unique transcriptional signature that is maintained long after the acute inflammation has resolved, suggesting permanent epithelial cell changes.^3^ Epigenetic changes in genes from UC mucosa have been suggested related to pathways that affect antigen processing and presentation, cell adhesion, B- and T-cell receptor signaling, JAK-STAT signaling, and transforming growth factor beta (TGF-β) signaling etc. However, the extent and consequences of epigenetic changes in IBD have not been adequately characterized. However, given that abnormal barrier function and a reduced protective mucin layer contribute to initiation of IBD and potentially to recurrence, the presence of both characteristics in colonoids over multiple passages and in colonoids made from inactive as well as active UC tissue, suggests that UC mucosa is primed for recurrence even in the absence of inflammation. We speculate that an approach to reverse these changes in UC colonoids has the potential to prevent UC recurrence and potentially to prevent the proximal spread of UC, a concerning and unmet need in UC management.

## Acknowledgements

We thank the Hopkins Conte Basic and Translational Digestive Diseases Research Core Center Integrated Physiology and Translational Research Enhancement Cores for their contributions to obtaining and maintaining colonoids.

## Contributors

VS designed the concept, supervised the study, conducted experiments, analyzed data, and wrote the manuscript. KJ, JY, RL conducted experiments. SL, HY, recruited patients and/or collected samples. SL, JI, JF established colonoids, NZ supervised colonoid culture.

## Patient consent

Obtained.

## Ethics approval

Johns Hopkins Medicine IRB committee.

