## Supplementary method for "Chronic inflammation in ulcerative colitis causes long term changes in gobletcell function"

**SUPPLEMENTARY MATERIALS AND METHODS**

**Organoid culture and monolayer formation:** Colonoid cultures were established as originally described by Sat*o et al.*^1^ Human colonoid were maintained as cysts embedded in Matrigel (Corning #356231, USA) in 24-well plates and cultured in Wnt3A, Rspon and Noggin containing growth media or non-differentiated media (NDM).^2^ Medium was replaced with fresh NDM medium every other day. Studies were carried out on passages 6-40. For generation of monolayer, colonoids were fragmented in Organoid Harvesting Solution (Trevigen, USA) and multiple wells were pooled together and resuspended in NDM after centrifugation. Colonoid fragments (in 100 μl NDM) were added onto 0.4 μm pore transparent polyester (PET) membrane 24-well cell culture inserts (Transwell; Corning, USA or Millipore, USA) pre-coated with human collagen IV (30 μg/ml; Sigma-Aldrich, USA). NDM (600 μl) was added to the bottom, and the cultures incubated at 37 °C, 5% CO_2_. Monolayers were cultured in a 5% CO_2_ atmosphere at 37°C. Growth medium was supplemented with Y-27632 (10µM) and CHIR99021(10µM) during the first 2 days after seeding. Formation of colonoid monolayers was monitored by measurement of transepithelial electrical resistance (TEER). Once monolayers became confluent, expansion medium was replaced with differentiation medium that was made by substituting Wnt3A, R-spondin1, and SB202190 in the expansion medium with the base medium. Five days later, paired undifferentiated and differentiated enteroid monolayers were studied.

**Quantitative Real-Time Polymerase Chain Reaction:** Total RNA was extracted from 3D cultures using the PureLink RNA Mini Kit (Life Technologies) according to the manufacturer’s protocol. Complementary DNA was synthesized from 1 to 2 μg of RNA using SuperScript VILO Master Mix (Life Technologies). Quantitative real-time polymerase chain reaction (qRT-PCR) was performed using Power SYBR Green Master Mix (Life Technologies) on a QuantStudio 12K Flex real-time PCR system (Applied Biosystems, Foster City, CA). Each sample was run in triplicate, and 5 ng RNA-equivalent complementary DNA was used for each reaction. Commercially available primer pairs from OriGene technologies were used. The following primer pairs were used: *LGR5*: [HP207145](https://www.origene.com/catalog/gene-expression/qpcr-primer-pairs/hp207145/lgr5-human-qpcr-primer-pair-nm_003667), *Ki67*: [HP206104](https://www.origene.com/catalog/gene-expression/qpcr-primer-pairs/hp206104/ki67-mki67-human-qpcr-primer-pair-nm_002417), *ATOH1*: [HP208359](https://www.origene.com/catalog/gene-expression/qpcr-primer-pairs/hp208359/atoh1-human-qpcr-primer-pair-nm_005172), *NEUROG3*: [HP213982](https://www.origene.com/catalog/gene-expression/qpcr-primer-pairs/hp213982/neurogenin3-neurog3-human-qpcr-primer-pair-nm_020999), *SPDEF*: [HP210328](https://www.origene.com/catalog/gene-expression/qpcr-primer-pairs/hp210328/spdef-human-qpcr-primer-pair-nm_012391), *MUC2*: [HP206138](https://www.origene.com/catalog/gene-expression/qpcr-primer-pairs/hp206138/muc2-human-qpcr-primer-pair-nm_002457), *CHGA*: [HP205193](https://www.origene.com/catalog/gene-expression/qpcr-primer-pairs/hp205193/chromogranin-a-chga-human-qpcr-primer-pair-nm_001275), *LYZ*: [HP200222](https://www.origene.com/catalog/gene-expression/qpcr-primer-pairs/hp200222/lysozyme-lyz-human-qpcr-primer-pair-nm_000239), *NHE3*: [HP207529](https://www.origene.com/catalog/gene-expression/qpcr-primer-pairs/hp207529/slc9a3-human-qpcr-primer-pair-nm_004174), *DRA*: [HP200096](https://www.origene.com/catalog/gene-expression/qpcr-primer-pairs/hp200096/slc26a3-human-qpcr-primer-pair-nm_000111), *CAII*: [HP200053](https://www.origene.com/catalog/gene-expression/qpcr-primer-pairs/hp200053/carbonic-anhydrase-ii-ca2-human-qpcr-primer-pair-nm_000067), *CFTR*: [HP200464](https://www.origene.com/catalog/gene-expression/qpcr-primer-pairs/hp200464/cftr-human-qpcr-primer-pair-nm_000492), *PAT1*: [HP232409](https://www.origene.com/catalog/gene-expression/qpcr-primer-pairs/hp232409/slc26a6-human-qpcr-primer-pair-nm_022911), *NHE1*: [HP206641](https://www.origene.com/catalog/gene-expression/qpcr-primer-pairs/hp206641/slc9a1-human-qpcr-primer-pair-nm_003047), *NKCC1*: [HP203742](https://www.origene.com/catalog/gene-expression/qpcr-primer-pairs/hp203742/nkcc1-slc12a2-human-qpcr-primer-pair-nm_001046), *KCNE3*: [HP208601](https://www.origene.com/catalog/gene-expression/qpcr-primer-pairs/hp208601/kcne3-human-qpcr-primer-pair-nm_005472), *NBCE1*: [HP232301](https://www.origene.com/catalog/gene-expression/qpcr-primer-pairs/hp232301/slc4a4-human-qpcr-primer-pair-nm_003759), *AE2*: [HP206636](https://www.origene.com/catalog/gene-expression/qpcr-primer-pairs/hp206636/ae2-slc4a2-human-qpcr-primer-pair-nm_003040), *NHE2*: [HP206642](https://www.origene.com/catalog/gene-expression/qpcr-primer-pairs/hp206642/slc9a2-human-qpcr-primer-pair-nm_003048), *18S rRNA*: [HP220445](https://www.origene.com/catalog/gene-expression/qpcr-primer-pairs/hp220445/rna18sn5-human-qpcr-primer-pair-nr-003286) (OriGene Technologies, Rockville). The relative fold changes in mRNA levels of genes between differentiated organoids and undifferentiated organoids were determined using the 2^-ΔΔCT^ method with human 18S ribosomal RNA simultaneously studied and used as the internal control for normalization and shown in fold‐change compared to the HS-UD or HS-DF control.

### Immunofluorescence staining, Confocal and TEM Image analysis

Analysis of MUC2 by immunofluorescence and confocal microscopy was carried out as previously reported.^3^ Briefly, human colonoid monolayers were fixed with Carnoy’s solution (90% [v/v] methanol, 10% [v/v] glacial acetic acid), washed three times with PBS, permeabilized with 0.1% saponin, and blocked with 2% bovine serum albumin+15% fetal bovine serum for 60 minutes (all Sigma-Aldrich, USA) followed by overnight incubation with antibodies. For immunostaining in 3D organoids, staining was done in suspension. Briefly, recovered organoids were fixed in 4% paraformaldehyde in 10mM phosphate buffer (pH 7.4) for 30 minutes at 4^0^C, then washed 2x with PBS. Organoids were permeabilized and stained in phosphate-buffered saline with 2% bovine serum albumin, 1% Triton X-100, and 1% saponin. Images were collected using a 20x or 40x oil immersion objective on an FV3000 confocal microscope (Olympus) with software (Olympus) and ImageJ software (NIH). Images were 3D-reconstructed using Volocity Image Analysis software (Improvision). Primary antibodies included Rabbit anti-MUC2 (Santa Cruz Biotechnology, USA; Cat#sc7314) and Rabbit anti-ChgA (Santa Cruz Biotechnology, USA; Cat#sc1488). All antibodies were diluted 1:100. For quantitative analysis, the same settings were used to image across samples (e.g. MUC2 staining). Mucin exocytosis and thickness was determined by measuring MUC2 positive area above the epithelial surface. For electron microscopy, 2-mm sections were fixed in 1% osmium tetroxide and 1% uranyl acetate, dehydrated with ethanol, and infiltrated with epoxy resin. Thin sections (80 nm) were cut and transferred to 200-mesh copper grids before staining with uranyl acetate and lead citrate. Grids were viewed on a Hitachi 7600 434 TEM operating at 80 kV and digital images of the apical regions were captured with an AMT 1K × 1K CCD camera.

***Escherichia coli* strains (ETEC) and infections**. ETEC H10407 strain was described previously. All antibiotics were purchased from Sigma Chemical Co. (St. Louis, MO). For colonoid infections, strain was grown from frozen stocks (−80 °C) at 37 °C on Luria broth (LB) agar plates (Difco) 2- days prior to experiments. Day before infection, single colonies were inoculated in 5ml of L-broth and grown overnight with vigorous shaking at 37 °C. For infection, an overnight LB culture was diluted 1:50 into fresh LB and incubated at 37°C with agitation for 2 h to achieve a log-phase culture (OD600=0.6). Subsequently, bacteria were adjusted to 10^8^ CFU/ml in sterile PBS, and 10μL (1×10^6^) was added to the apical surface of colonoid monolayers. *E. coli* infections were allowed to progress for 8h.
